# Frequency-dependent cerebellar circuits independently gate social vocalizations and movement

**DOI:** 10.64898/2026.02.18.706564

**Authors:** Cheryl Brandenburg, Snigdha Srivastava, Alejandro G. Rey Hipolito, Tao Lin, Benjamin R. Arenkiel, Roy V. Sillitoe

**Affiliations:** Department of Pathology & Immunology, Baylor College of Medicine; Houston, 77030, USA; Cerebellum Science Center, Texas Children’s Hospital; Houston, 77030, USA; Jan and Dan Duncan Neurological Research Institute at Texas Children’s Hospital; Houston, 77030, USA; Department of Molecular and Human Genetics, Baylor College of Medicine; Houston; 77030, USA; Medical Scientist Training Program, Baylor College of Medicine; Houston; 77030, USA; Department of Neuroscience, Baylor College of Medicine; Houston; 77030, USA; Department of Pediatrics, Baylor College of Medicine; Houston, 77030, USA

## Abstract

Communication depends on precise coordination between motor execution and cognition. Here we reveal that the cerebellum exerts real-time control over social vocalizations in adult mice. Optogenetic activation of excitatory cerebellar output in the superior cerebellar peduncle suppressed ultrasonic vocalizations with frequency-dependent potency while inducing distinct motor phenotypes. Systematic, functional mapping across cerebellar regions and cell types revealed that vocal suppression can occur in the absence of overt motor impairment, suggesting selective control of vocal output beyond gross movement disruption. We identified the periaqueductal gray (PAG), a conserved midbrain vocal control center, as a key downstream mediator of this effect. Deep brain stimulation of the PAG rescued vocal deficits in a model of cerebellar dysfunction without rescuing motor incoordination. These findings define a cerebellar–midbrain pathway that gates vocal behavior and demonstrate that targeted therapeutic neuromodulation can selectively restore communication-related output even in the presence of persistent cerebellar-driven motor deficits.

---

Language acquisition is a defining feature of human development, linking cognition to social communication. The neural circuits that govern language span distributed cortical networks (*1*, *2*), with the cerebellum often being overlooked. However, cerebellar participation in language (*3–8*) and other cognitive domains (*9–12*) has been increasingly recognized. As a critical hub that integrates motor and non-motor functions, cerebellar connectivity with the cerebral cortex dynamically adapts across development (*13*, *14*) in parallel with the emergence of higher-order cognition and coordinated motor behavior. Human imaging studies underscore the complex relationship between cerebellar circuits and behavior across functional domains (*15*, *16*), raising the question of how specific cerebellar regions contribute to distinct behavioral processes.

Many neurodevelopmental and neurodegenerative conditions that may be rooted in cerebellar dysfunction present with co-occurring motor and non-motor challenges that profoundly affect quality of life (*17*). Autism spectrum disorder (ASD), for example, is characterized primarily by differences in social communication, alongside variable motor and repetitive behavioral features, with converging evidence implicating cerebellar involvement (*18–24*). Despite this association, it remains unclear whether cerebellar circuits that influence communication are separable from those that control movement, or whether non-motor deficits arise secondarily from disrupted motor output. Parsing cerebellar circuits that differentially influence movement and communication is therefore a critical step toward uncovering mechanisms of cognitive dysfunction and identifying therapeutic targets. Animal models provide a powerful platform to address this problem using temporally precise circuit-level manipulations.

Ultrasonic vocalizations (USVs) in mice provide a sensitive and quantifiable readout of vocal output during social interaction and are widely used as a proxy for social communication. Whether cerebellar circuits directly influence ongoing adult vocalizations, however, has not been established. To address this, we manipulated cerebellar output at its primary efferent pathway to determine whether vocal behavior could be acutely modulated. We find that cerebellar output rapidly and reversibly suppresses USVs in adult mice, demonstrating that the cerebellum can directly influence ongoing vocal behavior. This suppression can be mediated through projections to the periaqueductal gray (PAG), a conserved midbrain vocal control center. We then systematically mapped cerebellar subregions and cell types using frequency-specific optogenetic stimulation, revealing that modulation of vocal output does not uniformly track with overt motor phenotypes, including tremor and motor incoordination.

## Cerebellar excitatory output gates vocal communication in mice

The immense computational processing that occurs within the cerebellar cortex may be distilled into its main output neurons. In the rodent, billions of granule cells make hundreds of thousands of synapses onto individual Purkinje cells, which are additionally contacted by several types of interneurons and one or two functionally potent climbing fibers. This extensive convergence enables Purkinje cells to integrate diverse synaptic inputs and translate them into a unified output signal. As the sole output neurons of the cerebellar cortex, Purkinje cells transmit this integrated activity via their axons (with local collaterals) predominantly to the cerebellar nuclei. Similarly, the main output pathway of the entire cerebellum is contained within the superior cerebellar peduncle (SCP), a white matter tract formed by the projections of cerebellar nuclei neurons that target multiple extra-cerebellar brain regions (*25*).

We capitalized on this organized output architecture by expressing channelrhodopsin (ChR2) in excitatory output cells of the cerebellar nuclei, enabling us to directly manipulate cerebellar signaling, and determine whether it influenced adult rodent vocalizations in a social behavior paradigm. As *Atoh1* is expressed in excitatory neurons of the cerebellum, including granule cells and nuclei neurons (*26*, *27*), we used it to drive Cre-dependent ChR2 expression. 3D light sheet imaging confirmed strong expression of *Atoh1*-tdTomato in excitatory cells of the cerebellar cortex (Fig. 1A, Movie S1). To selectively target cerebellar output, optic fibers were placed bilaterally over the SCP after it exits the cerebellar nuclei. This white matter tract contains the axons of cerebellar nuclei neurons and is spatially segregated from the cerebellar cortex by hundreds of microns (Movie S1). As a result, optical stimulation at this location only engages ChR2-expressing output axons. This strategy enables selective manipulation of cerebellar output pathways despite broader ChR2 expression in the cortex.

**Figure 1.**
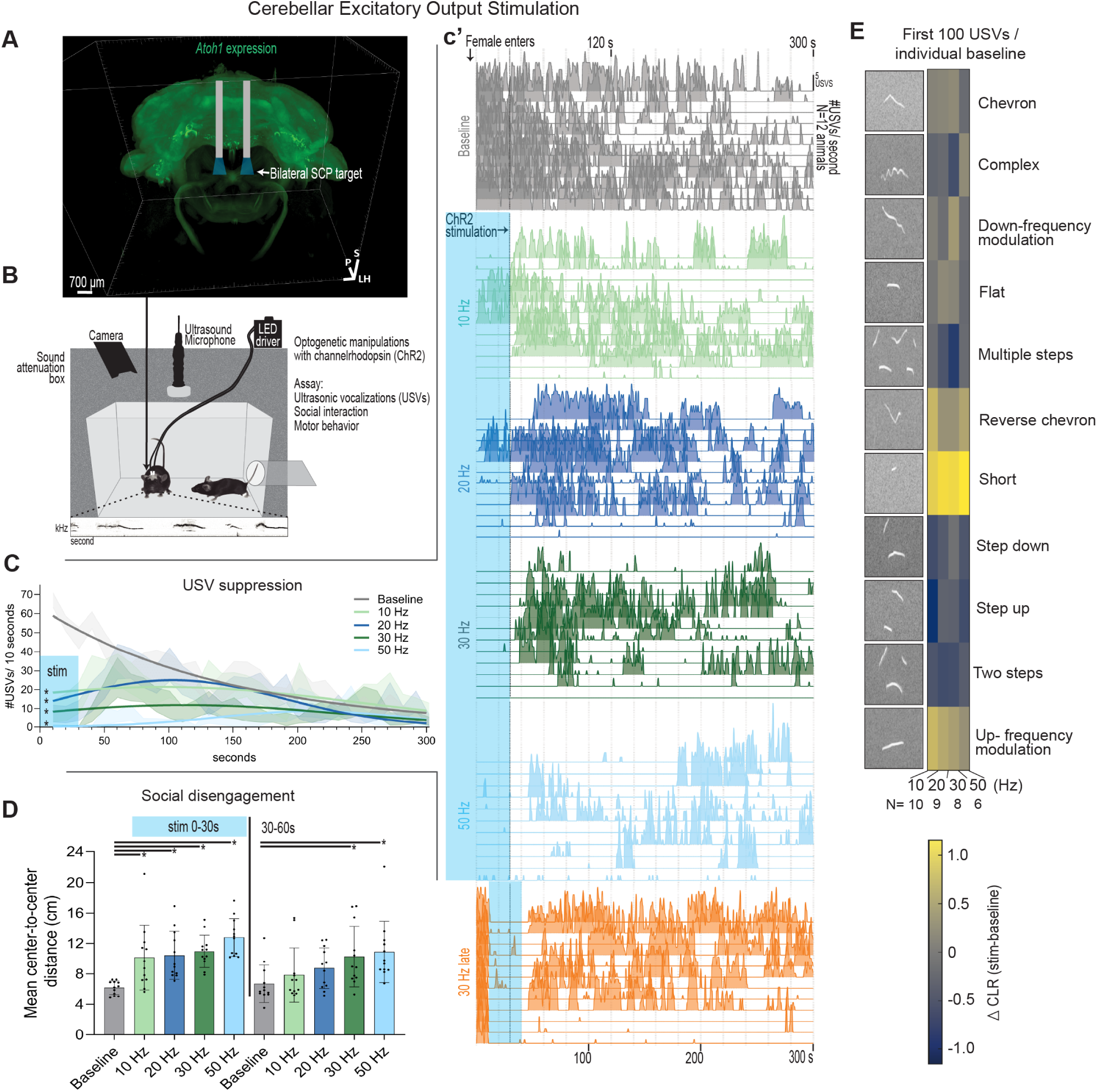
Cerebellar excitatory output via the superior cerebellar peduncle acutely suppresses ultrasonic vocalizations. **(A)** Light-sheet reconstruction of *Atoh1*-tdTomato expressing cerebellar excitatory neurons and their axonal projections. Bilateral optical fibers are positioned above the superior cerebellar peduncle (SCP) to selectively target cerebellar excitatory output pathways while avoiding stimulation of granule cells, Purkinje cells or inhibitory nuclei cells (Movie S1). **(B)** Behavioral assay schematic. Male mice implanted bilaterally above the SCP are recorded in a sound attenuated chamber during social interaction with a female. Ultrasonic vocalizations (USVs) are detected using an ultrasound microphone positioned above the chamber, synchronized with video tracking. Optogenetic stimulation consists of 10 ms square-wave light pulses delivered at 10, 20, 30 or 50 Hz during defined behavioral epochs. Representative USV traces are shown below the chamber schematic. **(C)** Time course of USV production aligned to female entry. Optogenetic stimulation of SCP excitatory output (blue shading) produces a rapid, frequency-dependent suppression of USVs. Quantification of the number of USVs produced during the first 30 s of stimulation revealed a significant effect of stimulation condition (one-way ANOVA, *F*(6,77) = 25.64, *p* < 0.0001). Dunnett’s multiple comparisons test showed that each stimulation frequency differed significantly from baseline (10 Hz: *p* < 0.0001; 20 Hz: *p* < 0.0001; 30 Hz: *p* < 0.0001; 50 Hz: *p* < 0.0001). Higher stimulation frequencies resulted in near-complete vocal silencing that often persisted beyond stimulation offset. (**c’**) Ridge plots depict individual animals (the same line across trials) tracked across stimulation frequencies. When stimulation is initiated 10 s after female entry, USVs are abruptly silenced and remain suppressed in a subset of animals following termination of stimulation. Each line represents the same animal across frequencies on separate days. *N* = 12 animals. **(D)** Social interaction quantified by the distance between the centers (torsos, Movie S3) of interacting animals during the first 0–30 s of stimulation and the subsequent 30–60 s interval. During the 0–30 s stimulation window, inter-animal distance differed significantly across conditions (one-way ANOVA, *F*(4,55) = 8.903, *p* < 0.0001). Dunnett’s post hoc comparisons revealed significant increases in distance relative to baseline at all stimulation frequencies (10 Hz: *p* = 0.0038; 20 Hz: *p* = 0.0018; 30 Hz: *p* = 0.0004; 50 Hz: *p* < 0.0001). During the 30–60 s post-stimulation interval, a smaller but significant effect of stimulation condition persisted (one-way ANOVA, *F*(4,55) = 3.042, *p* = 0.0245), with Dunnett’s comparisons indicating continued differences from baseline (30 Hz: *p* = 0.044; 50 Hz: *p* = 0.014). Error bars indicate mean ± SD; individual points represent animals. Additional comparisons shown in Fig. S1. *N* = 12 animals. **(E)** Composition of ultrasonic call types classified using VocalMat (27). Heatmaps depict the centered log-ratio (CLR) difference between stimulation and baseline conditions for the first 100 calls following vocal onset in each session. CLR values were computed per animal and averaged across animals within each stimulation frequency (10–50 Hz). Only animals producing ≥100 calls in both baseline and stimulation sessions were included (*N* = 10, 9, 8, and 6 animals for 10, 20, 30, and 50 Hz, respectively). No call type reached significance after false discovery rate correction when assessed using paired-sample effect sizes across animals; however, stimulation consistently biased vocal output toward short calls, indicative of a restricted vocal repertoire following SCP activation. Corresponding control animal distributions are shown in Fig. S1. Color scale denotes ΔCLR (stimulation − baseline).

For this experiment, male mice with bilateral SCP implants were habituated in a sound attenuation box containing an ultrasound microphone and camera to monitor USVs, social interactions, and motor behavior (Fig. 1B). In this behavioral paradigm, the males drive USVs, whereas females rarely vocalize (*28–30*). We focus on males due to their reliability to vocalize at specific time points that can be aligned with optogenetic stimulation during a social behavior. At baseline (Fig. 1C, gray), all 12 males (previously exposed to a female), began USV emissions after entry of a female into the chamber (without any before). Over the course of five minutes, USVs gradually tapered off, making the initial 30 seconds following female entry the period with the highest number and most reliable period for USV generation. We therefore chose to drive excitatory cerebellar output stimulation for 30 seconds preceding, and 30 seconds after female entry, when males are most motivated to vocalize. Increasing stimulation frequencies (10, 20, 30, and 50 Hz) showed higher likelihood to silence USVs (Fig. 1C, c’), with persistent silencing at higher frequencies, even after stimulation was ceased. Twelve control mice without ChR2 expression maintained USVs across each trial (Fig. S1). Each stimulation frequency and behavioral assay were performed on separate days using novel females to reduce variability of USV emissions. Each line in Fig. 1c’ represents the same male across trials.

To ensure that daily stimulation of the entire cerebellar excitatory output did not damage the ability of mice to generate appropriate USVs, and to determine the time-course of USV silencing effects, we next delayed the stimulation until 10 seconds after the female entered the chamber and observed that all males again increased USV number (Fig. 1 c’, orange). Immediately upon 30 Hz light pulse delivery USVs were robustly silenced throughout the 30 second stimulation, and the stimulation was sufficient to silence USVs for the duration of the assay in several mice. The observed silencing effect was not lateralized, as restricting the stimulation to either side could drive the effect (Fig. S1). Further, social interactions paralleled the USV changes, in that mice at baseline stayed very close together (measured by center-to-center distance, additional analyses in Fig. S1), while mice spent further time away from each other at increasing stimulation intensities (Fig. 1D). This effect manifested more intensely with higher stimulation frequencies, with males primarily avoiding females, while females explored freely (Movie S2). When USVs (first 100 emissions) resumed, we noted a tendency toward more up-frequency modulation and shorter calls (Fig. 1E). Overall, these data show that targeted activation of cerebellar excitatory output silences USVs and leads to decreased sociability in male mice.

## Functional mapping of segregated vocal and motor domains

Targeting excitatory projections within the SCP robustly suppressed USVs across stimulation frequencies; however, distinct and frequency-dependent motor phenotypes emerged in parallel. At 10 Hz, stimulation produced a prominent body-wide tremor. At 20 Hz, no overt motor phenotype was apparent, although a low-power tremor was detectable using an accelerometer-based tremor monitor (Fig. S2). At 30 Hz, neither tremor nor gross motor abnormalities were observed, whereas at 50 Hz, mild motor incoordination emerged. Stimulation at all frequencies increased inter-animal distance (Fig. 1D, Movie S2).

Because the SCP contains axons that originate from multiple cerebellar nuclei, it was unclear whether the specific circuits responsible for USV suppression were the same as those producing motor phenotypes. To disentangle these effects, we constructed a “phenotypic” heat map (Fig. 2) that summarized the presence or absence of vocal, social, and motor phenotypes across stimulation frequency, cerebellar region, and cell type. This structure-function mapping revealed a dissociation between vocal suppression and motor dysfunction. Notably, a robust 10 Hz tremor was elicited by stimulation of every cerebellar region tested and across excitatory neurons, inhibitory SCP projection neurons, and Purkinje cells, revealing that tremor is not strictly localized to a particular region or cell type. However, only a subset of these manipulations affected USVs. For example, stimulation of excitatory neurons in the dentate nucleus and/or Purkinje cell projections targeting the dentate nucleus reliably induced tremor, yet animals continued to generate appropriate USVs (Fig. 2, Fig. S3) and maintained social distance. Stimulation of SCP excitatory projections suppressed USVs and increased social distance when tremor was present, but stimulation of SCP inhibitory projections caused tremor with no effect on USVs or social interaction. Therefore, tremor alone was insufficient to account for deficits in social communication and vocal output, likely reflecting the downstream impact of these different pathways on particular regions.

**Figure 2.**
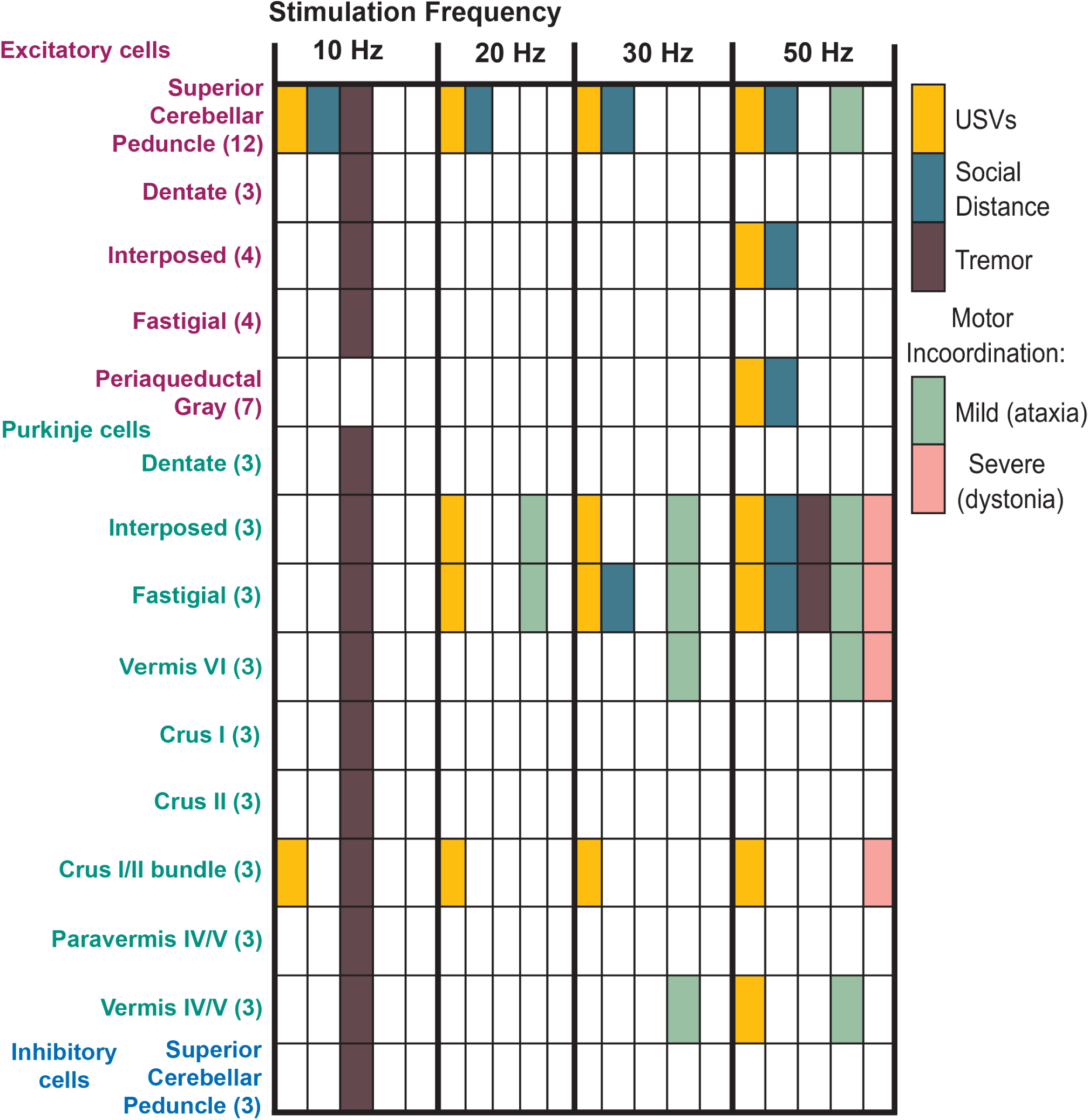
Frequency-specific functional mapping of social and motor phenotypes across cerebellar cells and regions. Binary summary of behavioral phenotypes elicited by optogenetic stimulation of distinct cerebellar subregions and cell types at 10, 20, 30, or 50 Hz. Phenotypic outcomes varied systematically with stimulation frequency, cell type, and anatomical target, revealing a frequency-dependent functional organization of cerebellar signaling. Ultrasonic vocalizations (USVs) were quantified using the machine-learning–based classifier VocalMat. Given the exploratory nature of this dataset and limited sample sizes for several stimulation targets, phenotypes were operationally defined using within-group consistency criteria rather than population-level statistical inference. Specifically, for groups with N < 5, a USV phenotype was scored as present if all animals exhibited a ≥20% reduction relative to baseline during stimulation (Fig. S3). Social disengagement was assessed by changes in center-to-center distance (Movie S3), with a phenotype assigned when all animals in a group showed a ≥12% increase in distance (Fig. S3). Superior cerebellar peduncle comparisons are shown in Fig. 1A and periaqueductal gray comparisons are shown in Fig. 3E. Tremor was identified through direct behavioral observation and independently confirmed using an accelerometer-based tremor monitor (Fig. S2) and marked positive when physiological tremor (increased power near 10 Hz) was present. Motor incoordination was classified as mild (ataxia) or severe (dystonia) based on overt abnormalities in posture and movement evident in behavioral videos (Movie S4). Severe motor incoordination co-occurred with increased social distance, consistent with impaired motor control contributing to apparent social disengagement under these conditions. Colored boxes indicate the presence of a phenotype under the specified stimulation condition, whereas white boxes indicate its absence. Numbers in parentheses denote the number of animals tested for each stimulation target. Quantitative analyses underlying each behavioral measure are provided in the supplementary figures. Bilateral optical stimulation was performed in *Atoh1*^Cre^ (excitatory output neurons and axon terminals in the PAG), *Pcp2*^Cre^ (Purkinje cells), and *Ptf1a*^Cre^ (inhibitory output neurons) mice crossed to *Rosa26*^lsl-ChR2-eYFP^. The PAG was targeted either bilaterally or unilaterally.

Further arguing against motor dysfunction alone physically constraining vocal output, severe motor incoordination that resembled dystonia could also co-occur with USV suppression (Crus I/II Purkinje axon bundle), or with normal vocalizations (vermis VI). Frequencies of at least 50 Hz were required to produce severe motor incoordination resembling dystonia, while ataxia could be produced with stimulation as low as 20 Hz when targeting Purkinje cells to the fastigial or interposed nuclei. Additionally, within the fastigial and interposed, a complicated mixture of motor phenotypes was apparent (Movie S3), with co-occurring dystonia, ataxia, and tremor similar to our previous findings (*32*). CrusI and CrusII are heavily implicated in cerebellar associations to language (*8*, *33*) and targeting the CrusI/II projection bundle en route to the nuclei led to silencing of USVs, while individually targeting subsets of cells within the lobules of Crus I or Crus II did not. The lateral areas of lobule IV/V (in the paravermis) did not silence USVs, but targeting the same lobule (IV/V) in the vermis reduced USVs during motor incoordination episodes. In some regions (Crus I/II bundle), 20 Hz did not cause motor incoordination, but a USV deficit persisted. These findings demonstrate that cerebellar suppression of vocalizations cannot be explained by gross motor impairment alone, and instead implicate downstream circuits capable of selectively gating communication.

## Cerebellar-periaqueductal gray circuit drives vocal suppression

We next considered regions with significant cerebellar projections that may contribute to USVs. The SCP projects widely throughout the brain, but we focused on areas likely to mediate USVs independent of gross motor disruption. Of the potential targets, the periaqueductal gray (PAG) was chosen due to its well-known role as a vocal control center in mammals (*34–36*), which is densely innervated by the cerebellum (*25*). Targeting the PAG may eliminate the optogenetically induced motor symptoms if it is specialized for cognitive/affective states or more specific vocal control mechanisms. Indeed, targeting cerebellar excitatory terminals within the PAG led to a substantial reduction in USVs with no apparent motor phenotypes (Fig. 2, Fig. 3A). Interestingly, the PAG was the only region targeted that did not display a tremor at 10 Hz (Fig. S2), suggesting that the PAG is a significant target of cerebellar output influencing USVs and not gross motor dysfunction. Both the SCP and the PAG stimulated animals had similar overall locomotion compared to control (no ChR2 expression) animals (Fig. S4).Optogenetic manipulations of cerebellar projections within the PAG led to a similar reduction seen in a previous study that directly manipulated PAG neurons (*36*). Therefore, projections from the cerebellum are sufficient to drive a reduction in USVs that resembles direct PAG modulation.

**Figure 3.**
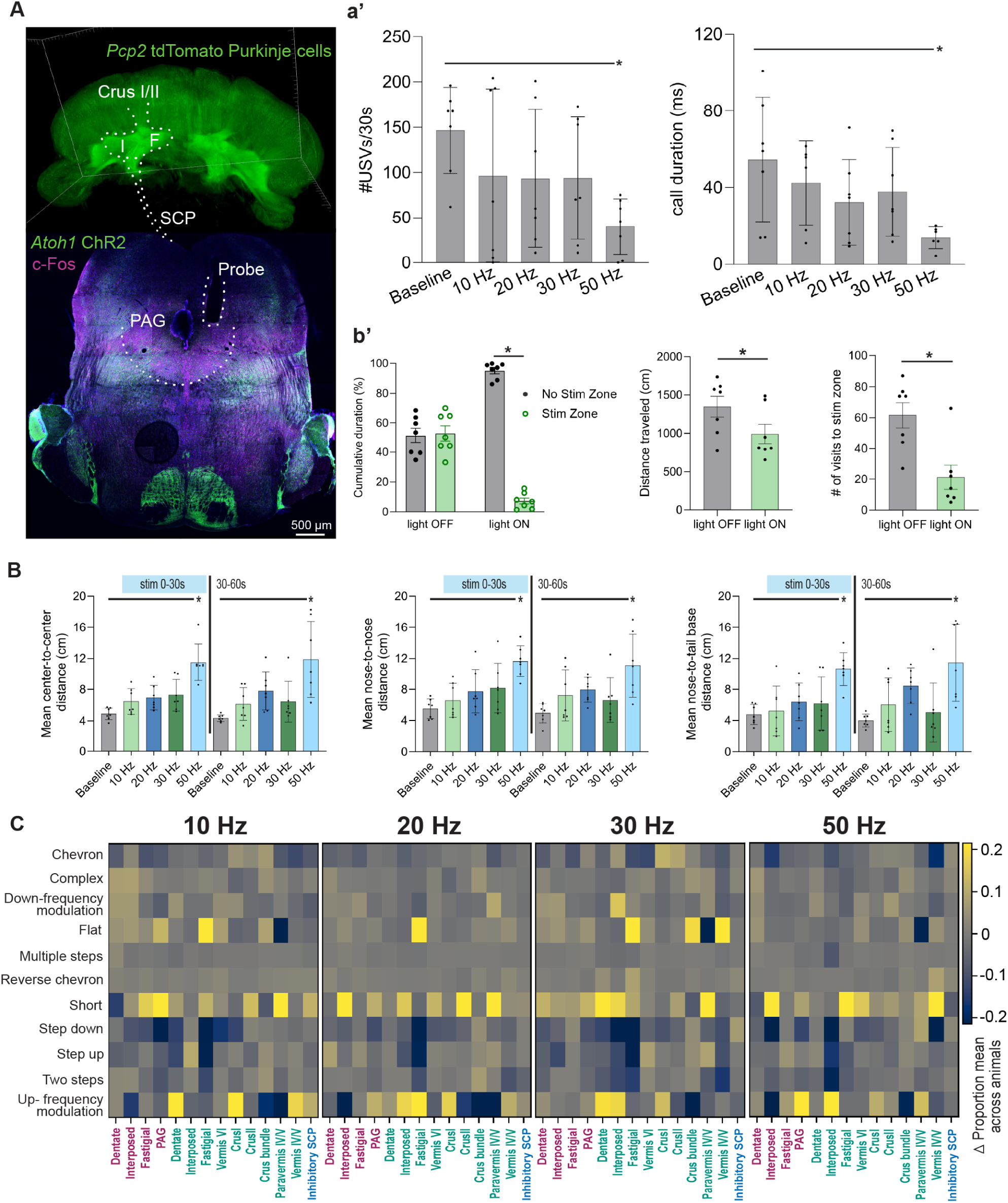
Cerebellar vocalization circuit map identifies a cerebellar–periaqueductal gray pathway that suppresses vocalization and encodes negative valence. **(A)** Summary circuit map of cerebellar regions where stimulation significantly suppressed ultrasonic vocalizations. Regions are indicated based on optogenetic targeting and behavioral outcomes summarized across experiments (see Fig. 2). Representative histological validation shows optic fiber placement and channelrhodopsin expression (green) in cerebellar projections terminating in the periaqueductal gray (PAG), along with induction of c-Fos (magenta) following PAG stimulation, confirming downstream activation of PAG neurons (Fig. S3) and general activity across the midbrain. I = interposed, F = fastigial, SCP = superior cerebellar peduncle, PAG = periaqueductal gray, Crus I/II = Purkinje cell fiber tract **(a’)** Direct stimulation of cerebellar excitatory projections within the PAG reduced vocal output. Optogenetic stimulation using laser pulses across frequencies decreased the number of ultrasonic vocalizations produced during the stimulation epoch (left) and reduced the duration of individual calls (right) relative to baseline. Each dot represents an individual animal; bars indicate mean ± SD. One-way ANOVA did not detect a significant main effect of stimulation frequency on USV number (*F*(4,30) = 2.184, *p* = 0.0948) or call duration (*F*(4,28) = 2.619, *p* = 0.0562). However, planned post hoc comparisons using Dunnett’s multiple comparisons test revealed a significant reduction at 50 Hz relative to baseline for USV number (adjusted *p* = 0.0209) and call duration (adjusted *p* = 0.0141). **(B)** Social interaction metrics during PAG stimulation. Mean center-to-center (left), nose-to-nose (middle), and male nose-to-female tail base (right) distances are shown for the stimulation epoch (0–30 s; blue bar) and the subsequent post-stimulation epoch (30–60 s). One-way ANOVA results 0-30 s: center-to-center *F*(4,30) = 13.86, *p* < 0.0001, 50 Hz Dunnett’s *p* < 0.0001; nose-to-nose *F*(4,30) = 5.539, *p* = 0.0018, 50 Hz Dunnett’s *p* = 0.0001; nose-to-tail base *F*(4,30) = 6.625, *p* = 0.0006, 50 Hz Dunnett’s *p* = 0.0008. 30-60 s: center-to-center *F*(4,30) = 13.86, *p* = 0.0005, 50 Hz Dunnett’s *p* = 0.0001; nose-to-nose *F*(4,30) = 4.510, *p* = 0.0057, 50 Hz Dunnett’s *p* = 0.0001; nose-to-tail base *F*(4,30) = 5.533, *p* = 0.0019, 50 Hz Dunnett’s *p* = 0.0010. Error bars indicate mean ± SD; individual points represent animals (*N* = 7). **(b’)** Real-time place preference (RTPP) assay demonstrates that activation of excitatory cerebellar projections to the PAG carries negative valence. When 50 Hz optogenetic stimulation was triggered upon entry into one side of the chamber, animals spent significantly less cumulative time in the stimulation-paired zone (left), traveled less total distance (middle), and made fewer entries into the stimulation zone (right) compared to light-off epochs. Gray bars denote non-stimulation periods; green bars denote stimulation periods. * indicates *p* < 0.05. **(C)** Call-type composition during optogenetic stimulation across cerebellar regions and stimulation frequencies. Heatmaps depict changes in call-type proportions during the first 30 s of stimulation relative to baseline. For each animal, the proportion of each call type was calculated during baseline and stimulation epochs, and the difference (Δ proportion) was summarized across animals. Call types are shown along the y-axis and cerebellar regions along the x-axis, grouped by stimulation frequency (10, 20, 30, and 50 Hz). No call-type category survived false discovery rate (FDR) correction across regions or frequencies; however, changes were directionally consistent across frequencies and regions, with a bias toward simpler call types, particularly short and up-frequency–modulated calls. Red region labels indicate excitatory cells, green are Purkinje cells and blue are inhibitory cells.

Having established the PAG as a key downstream mediator of vocal suppression, we summarize the overall regions that led to a silencing or reduction (Fig. 2, Fig. 3A). Bilateral optic implants to each cerebellar nuclei revealed that excitatory cells within the interposed nucleus were a primary driver of the silencing effect. In fact, neither the dentate nucleus nor the fastigial nucleus excitatory cells could drive the reduction. Since both excitatory cells of the interposed nucleus and the projections to those cells by Purkinje cells could silence USVs, this implicated the paravermis as a likely candidate, as the majority of interposed projections originate there. Most projections to the fastigial nucleus originate in the vermis, so lobules of the vermis may contribute as well (as seen with VI), with lateral lobules less likely since the dentate (lateral) nucleus did not drive USV changes targeting either excitatory nuclei cells or the Purkinje cell projections to them. The Crus I and II projection bundle could silence USVs at every frequency, as did the excitatory SCP output. The SCP excitatory projections terminating in the PAG reduced/silenced USVs and increased C-Fos across the midbrain after stimulation (quantification in Fig. S5).

Because PAG activation influenced both vocal output and social behavior, we next examined whether these effects reflected changes in affective state. When targeting both the SCP (Fig. 1D) and the PAG (Fig. 3B), males routinely disengaged from social interactions, measured by increased distance between the mice and was directly observable in the videos. The PAG has been shown to be involved in defensive/aversive behaviors (*37*) and projections from the cerebellum through the SCP are known to contribute to defensive behaviors driven by the PAG (*38*, *39*). Therefore, we tested whether, in addition to affecting USV properties (Fig. 3a’), activating cerebellar excitatory projections in the PAG was an aversive stimulus that could lead to the observed social disengagement, particularly because no motor impairments were observed. Using a real time place preference (RTPP) assay we found that when 50 Hz stimulation to the PAG was coupled to half of an arena, mice displayed a robust aversion to this stimulation and quickly avoided the stimulation side of the box (Movie S6), leading to less overall distance traveled and decreased entries to the stimulation side (Fig. 3b’). As the mice spent roughly equal time in each side of the box without light stimulation, these data indicate that cerebellar excitatory projections to the PAG can elicit negative valence, and may contribute to both avoidance behaviors during social interaction paradigms, and inter-animal distance (Fig. 3B).

Finally, stimulation of the PAG resulted in shorter duration calls (Fig. 1a’), which led us to expect that overall vocal repertoire could shift depending on the region manipulated. Indeed, cerebellar stimulation of specific cell types across multiple regions was found to induce a marked simplification of the vocal repertoire, defined by a relative enrichment of short and up-frequency modulation call types and a depletion of more complex, multi-step calls (Fig. 3C). Overall, we conclude the cerebellum gates USVs by eliciting negative valence as a state shift, which leads to silencing or simplifying vocal output.

## Neuromodulation restores vocalization deficits in a model with cerebellar circuit dysfunction

Collectively, our data show that altered cerebellar output to the PAG disrupts vocalizations and leads to an aversive state. Conversely, we wondered whether mice with severe cerebellar deficits could improve vocalizations with deep brain stimulation (DBS) targeted to the PAG. Our lab had previously shown that pup vocalizations were reduced when excitatory signaling is abolished from climbing fibers to Purkinje cells from birth (*40*), a model with persistent severe motor incoordination resembling dystonia (Fig. 4A-C) that could improve with DBS targeted to the interpose nucleus (*41*). For this, we first determined typical vocal behaviors across the lifespan by recording pup vocalizations in a maternal separation paradigm at postnatal day 0, 4, 7, 14, and 21 (Fig. 4b’). At P0, there was no significant difference between conditional knockout mice (CSKO, *Ptf1a*^cre^; *Vglut2*^fl/fl^) and their littermates, reflecting the feature that climbing fibers are just beginning to contact Purkinje cell somas with immature synapses. At P4, when climbing fibers are capable of eliciting EPSCs at Purkinje cell somas, USV reduction was apparent, though the pups could still produce USVs, which persisted at P7 (replicating our previous report). However, while control pups began to reduce the number of USVs generated, the CSKO mice showed more USVs at P14, potentially reflecting a delay in development as the climbing fibers move from soma to Purkinje cell dendrites to form their mature connections (*42*). By P21, both control and CSKO pups do not elicit USVs after maternal separation, showing that USV’s cease at an appropriate time point of development. This established that mice are physically capable of generating USVs during development and that the timeline of their disruption correlates with specific cerebellar functional circuit establishment.

**Figure 4.**
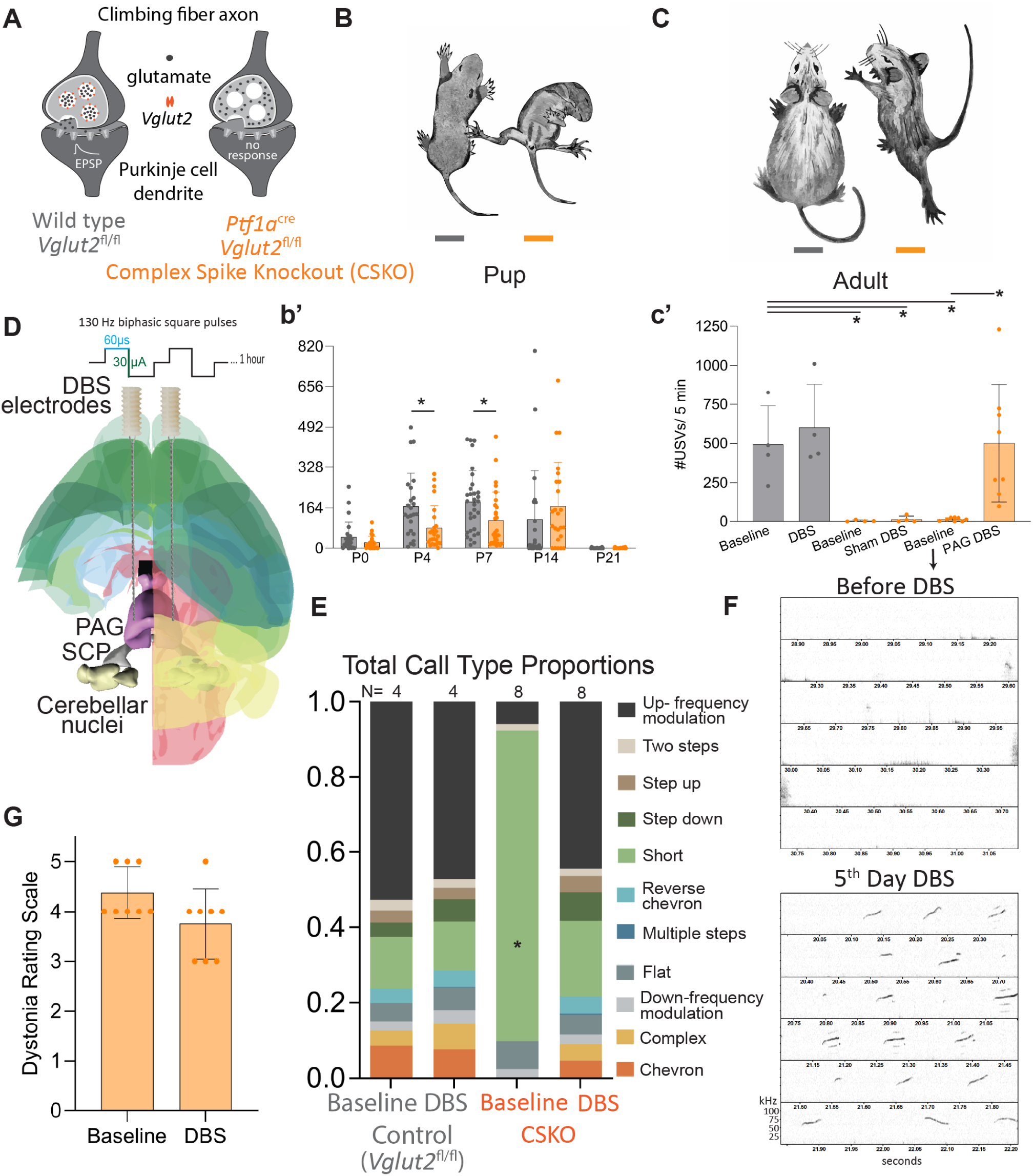
Neuromodulation of the periaqueductal gray (PAG) restores vocalization deficits in a complex spike knockout mouse model. **(A)** Schematic illustrating elimination of vesicular glutamate transporter 2 (*Vglut*2) selectively from the olivocerebellar pathway in *Ptf1a*^cre/+^; *Vglut*2^fl/fl^ mice (complex spike knockout; CSKO). Loss of *Vglut*2 abolishes fast glutamatergic neurotransmission from inferior olive climbing fibers onto Purkinje cells and cerebellar nuclei neurons, eliminating complex spike activity in Purkinje cells. **(B–C)** CSKO mice exhibit a pronounced dystonic motor phenotype beginning in early postnatal development (P7) that persists into adulthood. **(b’)** Developmental timeline of ultrasonic vocalizations (USVs) in dystonia pups. A two-way ANOVA revealed a significant interaction between age and genotype on pup ultrasonic vocalization (USV) production (*F*(4,242) = 3.076, *p* = 0.0170), indicating that the effect of complex spike knockout on vocal output depends on developmental stage. There was a significant main effect of age (*F*(4,242) = 15.76, *p* < 0.0001), reflecting robust developmental changes in USV production. In contrast, there was no significant main effect of genotype when averaged across ages (*F*(1,242) = 2.965, *p* = 0.0864), consistent with a developmentally restricted rather than global genotype effect. Post hoc simple-effects comparisons between control and dystonia pups at each age using Šídák correction did not identify any individual age at which the groups differed significantly (all adjusted *p* > 0.05). However, given the known developmental timeline of climbing fiber–Purkinje cell circuit establishment, we performed planned comparisons at specific early postnatal ages. These analyses revealed significant reductions in USV production in dystonia pups at P4 (*p* = 0.010, *N* = 24 control, 25 CSKO) and P7 (*p* = 0.020, *N* = 31 control, 28 CSKO) with unpaired t-tests. **(c’)** In adulthood, CSKO males exhibit a marked reduction in USV production during social interaction compared with control mice. Baseline USV output in CSKO males was significantly lower than that of control animals (unpaired t-test, *p* = 0.0077) and remained significantly reduced after sham DBS experiments with repeated exposure to novel females (*p* = 0.0085). *N* = 4 mice. In a separate cohort of males that showed reductions at baseline (*p* = 0.0002), deep brain stimulation (DBS) of the PAG significantly increased USV production in CSKO animals (paired t-test, *p* = 0.0080), restoring vocal output toward control levels. *N* = 8 mice. Gray bars indicate control mice; orange bars indicate CSKO mice. Points represent individual animals; bars show mean ± SD. **(D)** Schematic of deep brain stimulation targeting the periaqueductal gray (PAG). Biphasic stimulation was delivered at 130 Hz (60 µs pulse width, 30 µA) for 1 hour per day across 5 consecutive days. **(E)** Call-type composition analysis shows that short calls are disproportionately represented in CSKO mice at baseline, consistent with a restricted vocal repertoire. Following PAG DBS, call-type distributions shift toward those observed in control animals, driven primarily by a reduction in the proportion of short calls (paired t-test on short-call proportion, *p* = 0.0010). *N* = 4 control, 8 CSKO **(F)** Representative spectrograms from the same CSKO animal before DBS and after the fifth day of PAG DBS illustrate restoration of call diversity and increased vocal output. **(G)** Despite robust rescue of vocalization deficits, dystonic motor symptoms, quantified using a standardized dystonia rating scale, are not significantly improved by PAG DBS. Illustrations adapted from (42, 57).

We next tested whether severely dystonic CSKO mice have disrupted USVs in adulthood. To give the best chance of the male mice vocalizing, each CSKO mouse utilized was experienced, in that they produced at least one litter and were co-housed with a female for several weeks. They were then singly housed, implanted with DBS electrodes to the PAG, and allowed at least one week of recovery before the social interaction paradigm described in Fig. 1B was initiated. Baseline USVs of control vs CSKO adults were recorded over three days, and we report the third trial for each animal as the baseline number. CSKO mice exhibited limited vocalizations (Fig. 4c’), and this deficit was not improved by either sham DBS (no electrical stimulus delivered) or repeated exposure to novel females across five days. However, delivering 1 hour of DBS (130 Hz, 30 µA, 60 µs, Fig. 4D) to the PAG for five days was able to rescue the USV deficit (measured 30 minutes into the DBS protocol). Unlike our previous observation when targeting the interposed nucleus with DBS (*41*), dystonic motor symptoms were not rescued (Fig. 4G). The call types of CSKO adults were dominated by short calls before DBS, while after DBS the full range of typical call categories were observed (Fig. 4E-F).

Given that the main phenotype observed in the CSKO mouse is the elimination of complex spikes from Purkinje cells, this demonstrates that cerebellar signaling to downstream regions during postnatal and adolescent development may be important for driving social vocalizations in adulthood. Before climbing fiber circuitry is established (P0), pups generate typical USVs, suggesting that the cerebellum reshapes vocal behavior across development, which was recently demonstrated by assessing USV spectral properties in this dystonic model (*43*). Even though complex spikes are still eliminated in adulthood after DBS, which significantly disrupts cerebellar function and motor coordination, DBS to the downstream PAG could “bypass” the cerebellar dysfunction and promote USV generation in adults even with continued severe motor incoordination. This suggests that when cerebellar dysfunction is present, neuromodulation targeted to impactful downstream areas for specific behaviors may serve to rescue select behavioral deficits, and after critical periods of development.

## Discussion

We define a causal cerebellar–midbrain circuit that directly gates social communication, bridging a longstanding gap between motor control and higher-order behavior through the mapping of behaviors to specific cerebellar structures and cell types to distinct behavioral outputs. Optogenetic manipulation of excitatory cerebellar output robustly suppressed vocalizations, a finding with direct relevance to human clinical observations. In pediatric patients, surgical disruption of the SCP during posterior fossa tumor resection is frequently associated with cerebellar mutism syndrome, characterized by mutism, social withdrawal, and motor deficits (*44*). These features bear a striking resemblance to the phenotypes observed following SCP manipulation in mice. By identifying projections from the cerebellum to the PAG, we reveal a subcortical relay through which cerebellar output may influence affective and social behaviors. Consistent with this, recent imaging studies have linked cerebellar–PAG dysfunction to mutism in human patients (*45*), highlighting the overlapping functional capacities of this circuit across species.

Overall, low frequency stimulation reliably entrains motor loops, as a 10 Hz stimulation drove robust tremor across all cerebellar regions and cell types targeted. Intermediate frequencies (20 - 30 Hz) more clearly decouples motor vs affective circuits, and high frequency stimulation (50 Hz) tended to prevent vocal output and simultaneously cause severe motor incoordination. Combined, it appears that, under these conditions, the cerebellum does not encode vocal motor patterns per se, but rather regulates the permissiveness and timing of vocal output by modulating midbrain affective/vocal gating nodes (PAG), thereby biasing whether vocalizations are permitted, suppressed, or simplified.

Several observations support this interpretation. First, vocalizations were often preserved during the most severe motor disturbances, including tremor, ataxia and dystonia, depending on the region targeted. Second, animals maintained physical social interaction and produced USVs even under conditions of substantial motor disruption, arguing against a simple impairment of the vocal apparatus or gross motor impairment driving differences. Third, during stimulation conditions that suppressed vocal output, animals exhibited clear behavioral avoidance despite preserved locomotion, suggesting a shift in motivational or affective state rather than a purely motor deficit. Finally, direct stimulation of cerebellar projections within the PAG reduced, but did not abolish vocalizations and altered their acoustic features while preserving broad call categories, further indicating that vocal suppression is not explained by an inability to generate motor output.

Together, these findings indicate that cerebellar output can suppress vocal behavior independently of overt motor impairment. When vocalizations were reduced or absent, particularly in the PAG, this was accompanied by behavioral disengagement rather than generalized motor incapacity, supporting a model in which cerebellar signaling influences the decision to vocalize rather than the mechanics of vocal production within these behavioral paradigms. An important question that emerges is what information is conveyed by the cerebellar–PAG pathway. Real-time place preference assays suggest that activation of this pathway is aversive, consistent with growing evidence that cerebellar outputs engage PAG circuits involved in defensive and fear-related behaviors (*39*, *46*). These findings raise the possibility that cerebellar activity modulates vocal output by influencing internal state, biasing behavior away from social engagement. While USVs ultimately require motor execution, our data suggest that cerebellar contributions occur upstream of brainstem motor circuits, potentially regulating whether vocalization is initiated. Consistent with this, animals retained the capacity for locomotion but exhibited a shift from active pursuit to avoidance, supporting a reorganization of behavior rather than a loss of motor function. Although these results are best explained by a change in behavioral/brain state, we cannot exclude more subtle influences on motor control of the vocal apparatus.

Remarkably, DBS of the PAG restored vocal output in mice with genetically induced cerebellar dysfunction, even when severe dystonia persisted. This finding demonstrates that targeted neuromodulation can bypass impaired cerebellar computations to reinstate specific behavioral functions, a principle with broad translational potential for cerebellar ataxias, dystonias, and neurodevelopmental disorders that involve communication deficits. Determining the precise neural mechanisms that reinstate vocal output without restoring gross motor coordination will be important for determining how to leverage neuromodulation in promoting vocal output. Overall, disrupting key nodes in the vocalization network compresses output toward brief, simplified calls: dystonic mice exhibit short-call dominance that is reversed by DBS, PAG manipulation produces significantly shorter calls, and optogenetic stimulation across many regions shows a consistent trend toward short and up-frequency modulation calls, suggesting that overall disruption to cerebellar signaling biases vocal output toward a simpler repertoire.

Our work frames the cerebellum as an integrative hub that regulates the temporal precision of social signaling, linking motor, cognitive, and affective domains through overlapping, yet distinct circuitry. Frequency-specific activation patterns separately drive communication and movement behaviors, underscoring the importance of rate codes as behavioral gates. Functional mapping reveals dedicated locations where phenotypes originate, highlighting functional specificity at the level of the cerebellar cortex and nuclei driven by defined cell types. The restoration of vocal behavior through downstream stimulation further suggests that communication deficits resulting from cerebellar pathology may be therapeutically tractable even beyond developmental critical periods. Together, our results begin to illuminate how the cerebellum contributes to flexible coordination of complex behaviors, advancing a unifying framework for cerebellar function across domains of movement, affect, and communication.

## Acknowledgments

We sincerely thank all members of the Sillitoe lab for useful discussions and technical advice, particularly Dr. Amanda M. Brown, Dr. Linda H. Kim, Sarah Donofrio and Dr. Jason S. Gill.

## Funding

Simons Foundation SFI-AN-AR-Fellow-00012389 (CB)

Department of Defense PR22135 (BRA)

National Institute of Diabetes and Digestive and Kidney Diseases R01DK138518 (BRA)

National Institute of Diabetes and Digestive and Kidney Diseases R01DK109934 (BRA)

United States Department of Agriculture CRIS 1-58-3092-0-001 (BRA)

National Institute of Neurological Disorders and Stroke R01NS119301 (RVS)

National Institute of Neurological Disorders and Stroke R01NS127435 (RVS)

Eunice Kennedy Shriver National Institute of Child Health and Human Development of the National Institutes of Health P50HD103555 for use of the Cell and Tissue Pathogenesis Core (BCM IDDRC)

Baylor College of Medicine, Texas Children’s Hospital (RVS)

Ting Tsung and Wei Fong Chao Foundation (RVS)

## Author contributions

Conceptualization: CB, RVS

Methodology: CB, SS, AGRH, TL

Investigation: CB, SS

Visualization: CB, SS

Funding acquisition: CB, BRA, RVS

Supervision: BRA, RVS

Writing – original draft: CB

Writing – review & editing: CB, SS, AGRH, TL, BRA, RVS

## Competing interests

Authors declare that they have no competing interests.

## Data and materials availability

All data are available in the main text or the supplementary materials.

## Materials and Methods

### Animals

Mice were housed with a 20:00-6:00 dark cycle within a level 3, AALAS-certified vivarium. The Institutional Animal Care and Use Committee of Baylor College of Medicine reviewed and approved all experimental animal procedures under protocol AN-5996. The following transgenic mouse lines on a C57BL/6J background were utilized: *Atoh1^Cre^* (*47*), *Pcp2^Cre^* (also known as *L7^Cre^*) (*48*), *Ptf1a^Cre^* (gift from Dr. Chris Wright) (*49*), *Slc17a6^tm1Lowl^*/J (also known as *Vglut2^flox^;* JAX:012898) (*50*), *Ai32* (*Rosa26^lsl-ChR2-eYFP^*, JAX:012569) (*51*) and *Ai65* (*Rosa^lsl-tdTomato^*, JAX:007908) (*52*). The conditional knock-out mice that resulted in dystonia were generated by crossing *Ptf1a^Cre^;Vglut2^fx/fx^*with homozygote *Vglut2^fx/fx^* mice (*41*). Heterozygous expression of *Cre* in males was confirmed through ear punches and genotyping of each animal, then confirmed by positive expression of ChR2/tdTomato in each cell type when relevant. Due to the behavioral paradigm, we analyzed males (2-6 months old) only in this study, due to their reliability to produce USVs in response to a female and overrepresentation of developmental speech deficits in humans.

### Clearing and light sheet imaging

Whole brains from *Atoh1^Cre^*; *Rosa^lsl-tdTomato^*or *Pcp2^Cre^*; *Rosa^lsl-tdTomato^* were first cleared in Cubic-L at 37° for delipidation (*50–52*) for one-two weeks (changing solution every few days), rinsed with distilled water then placed in EZ Clear (*53*) for two days until imaging with a Zeiss Light Sheet Z1 at a refractive index of 1.52 with a 5x objective. Image tiles were stitched together with Stitchy and visualized with Imaris.

### Surgery

The same surgical technique was performed to implant 200 µm diameter optical fibers (typically bilateral) (ThorLabs, R-FOC-L200C-39NA) or two custom 75 mm twisted platinum DBS electrodes (PlasticsOne, Roanoke, VA, USA; 8IMS3039SPCE) into the various brain regions (Fig. S6) calculated through Pinpoint (*57*). Preemptive analgesics (buprenorphine, 1 mg/kg subcutaneous (SC), and meloxicam, 5 mg/kg SC) were delivered, with additional meloxicam doses the following 3 days as part of the standard post-operative analgesic protocol. Mice were continuously anesthetized with isoflurane while secured in a stereotaxic device (David Kopf Instruments), fur was removed with Nair, the skin was disinfected with alternating alcohol and betadine wipes, a midline incision was made and the skull was exposed and lightly etched with a blade. Small craniotomies were made at the target coordinates, the optical probes or DBS electrodes were slowly advanced into the brain, antibiotic ointment was added to cover the craniotomy, C&B Metabond (S380) was applied and allowed to harden then dental cement (A-M systems, #525000 and #526000) was applied over the top. Behavioral experiments started typically one week or more after the day of surgery. After behavioral experiments, brains were perfused, sectioned with a vibratome and targeting positions confirmed with fluorescent nissl staining and/or endogenous ChR2 expression on a Zeiss Axio Zoom. Only mice with confirmed appropriate expression and implant targeting (directly above the region of interest) were included in the dataset.

### Optogenetics and behavioral assay

Male mice with optical implants were attached to a bifurcated bundle (Thor Labs, BFYL2LS01) and fiber coupled LED (M470F4) connected to a Thor LED driver (LEDD1B) or Crystalaser (CL473) then allowed to habituate in the sound attenuation chamber for ten minutes. An Avisoft condenser ultrasound microphone (CM16/CMPA #40011) hovered 27 cm above the 27 cm wide Plexiglass animal chamber for ultrasonic vocalization (USV) recordings with UltraSoundGate 116B (#51161) and Avisoft-RECORDER USGH software. The Avisoft software was initiated simultaneously with Spike2 software for video behavior recordings and time-stamped delivery of stimulation pulses via the LED driver through a CED Power1401 data acquisition interface (CED, Cambridge, UK). All stimulation paradigms were delivered at full power (output 2.1 mW) with square pulses of 10 ms, with varying times between pulses (typically 10, 20, 30 or 50 Hz).

For the core behavioral paradigm used across all regions, baseline behavior of the male alone in the chamber was video recorded for one minute, the optogenetic stimulation paradigm was introduced and proceeded for 30 seconds, the female was introduced so that it fully entered the chamber through a side port by 90 seconds, then the optogenetic stimulation was terminated at 120 seconds (30 second stimulation alone, 30 second stimulation with female for a total of one minute stimulation) and the mice were allowed to freely interact for a total of 5 minutes. Variations to this protocol are indicated in figures/legends, such as when the stimulation was introduced after the female entered to abruptly terminate USVs when targeting the SCP (Fig. 3C orange). Each session was conducted on separate days with novel females, generally increasing frequency on subsequent days, but randomizing frequencies resulted in the same phenotype. Control males vocalized consistently across days with no effect from the stimulation (Fig. S1). Males did not vocalize when alone in the chamber (before female entry) and were singly housed before and between experiments.

### Deep Brain Stimulation

Male mice implanted with DBS electrodes (targeting the PAG; AP −4.7 mm, ML ±0.7 mm, DV −2.21 mm relative to bregma) were connected to the deep brain stimulation device (Multi Channel Systems, #STG4002) to deliver a biphasic 130 Hz stimulus with 60 µs duration at 30 µA for 1 h per day for 5 days (*41*). A female was introduced, as described in the optogenetic behavioral methods, with one full assay occurring 30 minutes into the one hour of DBS stimulation. Videos during the assay were examined for dystonic postures. All dystonic males were co-housed with females for several weeks and produced at least one litter before being single housed. Baseline USVs were recorded on the third trial (to allow the mice to have experience with the assay) while tethered to the DBS system, but no pulses were delivered (as with the sham DBS trials). Each trial was on a separate, consecutive day.

Dystonia rating scale (*55*) from 0 to 5 was used wherein 0 = no motor abnormalities; 1 = slightly slowed or abnormal motor behavior, no dystonia; 2 = mild impairment, sometimes limited ambulation, dystonic postures when disturbed; 3 = moderate impairment, frequent spontaneous dystonic postures; 4 = severe impairment, sustained dystonic postures and limited ambulation; and 5 = prolonged immobility in dystonic postures.

### Tremor, ataxia and dystonia behavior

Motor behavior was visually identified from videos taken during the behavioral assay (Movie S4) and binary presence of tremor, ataxia and dystonia were manually recorded by at least two expert observers. To confirm the presence of a tremor induced by the optogenetic stimulation and to verify the frequency of tremor induced with different stimulation paradigms, the mice were placed in a custom-made tremor monitor as described previously (*56*). Frequency and amplitude of the tremor were measured before, during and after optogenetic stimulation via optical patch cables connected above the tremor monitor, which is a Plexiglass box suspended by elastic chords at each corner with an accelerometer mounted to the box to detect subtle shifts in box movement. Mice were gently encouraged to ambulate a similar amount during each recording period for consistent baseline movement of the box. The output from the tremor monitor was amplified and lowpass filtered at 5 kHz (Brownlee Precision, Model 410) before being digitized by a CED board and analyzed using Spike2 Scripts. Recordings are centered on zero, downsampled using the Spike2 interpolate function and a power spectrum analysis with a Hanning window was performed for each recording window of 2 minutes (before, during and after stimulation). The peak power was calculated as the maximum power within the frequency band window. The mice are allowed to freely ambulate within the box, and given 2 minutes to acclimate before sequential 2 minute recording periods.

### Machine learning software analysis

USVs were recorded using Avisoft software and exported as .wav files for downstream analysis. USV files were processed in MATLAB (MathWorks) using VocalMat, an automated computer vision–based pipeline for detection and classification of ultrasonic calls (*57*). VocalMat identifies individual vocalizations based on spectral features and temporal structure then assigns calls to predefined categories using supervised machine learning classifiers trained on annotated datasets. Quantitative features describing call timing, duration, and spectral content were exported for each recording. Custom Python scripts were used to aggregate vocalization metrics across trials and animals, and summary data were visualized and statistically analyzed in GraphPad Prism (version 10.3.1 for Windows).

To quantify social interaction dynamics during male–female encounters, pose estimation was performed using Social LEAP Estimates Animal Poses (SLEAP) (*58*). A custom multi-animal bottom-up deep learning model was trained to detect and localize individual body landmarks (nose, ears, neck, mid-torso, tail base, tail midpoint, and tail tip) for each animal in video frames (Movie S3). The bottom-up architecture was selected because it provided robust performance across variable postures and partial occlusions typical of freely interacting animals. Model training produced confidence maps for individual body parts and part-affinity fields used to associate detected keypoints into distinct animal instances.

For inference, pose predictions were generated across entire video sequences using a multi-animal bottom-up inference pipeline with a maximum of two tracked instances. Identity tracking across frames was performed using optical-flow–based instance matching with greedy assignment, constrained by instance-level similarity metrics. Tracking incorporated a temporal window of five frames and optional Kalman filter–based smoothing after an initial warm-up period to improve cross-frame identity consistency. Single-frame track discontinuities were automatically reconnected during post-processing. Output pose trajectories were used for downstream quantification of movement metrics and between animal spatial relationships using custom Python analysis scripts.

### Real-time place preference assay

*Atoh1*-Cre mice expressing ChR2 with optic fibers targeting the periaqueductal gray (PAG) were assessed after USV assays. Both unilateral (n = 2) and bilateral (n = 5) fiber implant configurations were used. Mice were tested in a rectangular open-field arena divided into two spatially distinct halves. On the first day, animals were allowed to freely explore the arena for 30 minutes in the absence of photostimulation to establish baseline place preference. On a subsequent testing day, animals were returned to the same arena, and photostimulation (473 nm light, 50 Hz, 10 ms pulses) was delivered when the animal occupied one predefined half of the arena. The stimulated side was randomized across animals. No stimulation was delivered when animals occupied the opposite half of the arena.

Animal position and locomotor behavior were continuously tracked using EthoVision software (Noldus), which was used to quantify time spent in each half of the arena and to assess spatial occupancy. All behavioral metrics were computed from EthoVision outputs and visualized with Graph Pad Prism. Place preference was assessed by comparing the proportion of time spent in the stimulated versus unstimulated zones across sessions.

At the conclusion of the assay, mice were stimulated for 30 seconds on and 90 seconds off for 10 minutes at 50 Hz to induce c-Fos expression. Mice were perfused one hour later, brains were cryoprotected with 30% sucrose, sectioned on a cryostat and immunostained against GFP (Invitrogen A10262, 1:1000, Secondary: Invitrogen 11039 1:500) to detect ChR2 expression and c-Fos (Abcam ab190289, 1:1000, Secondary: Invitrogen A-21245 1:500). Sections containing the PAG implant were compared to control mice without stimulation to detect upregulation in c-Fos (Fig. S4).

### Statistics

All statistical analyses were performed using GraphPad Prism (version 10.3.1 for Windows) and custom Python scripts for data aggregation and preprocessing. No statistical methods were used to predetermine sample sizes; sample sizes were chosen based on prior studies using comparable behavioral and optogenetic paradigms. Data distribution was not formally tested for normality due to modest sample sizes; parametric tests were selected based on experimental design and robustness to minor deviations from normality. All tests were two-tailed and an alpha level of 0.05 was used to determine statistical significance. Exact *p* values, test statistics, and degrees of freedom are reported in figure legends where applicable.

#### Ultrasonic vocalizations

USVs were quantified as total call number within defined time windows (typically the first 30 s of stimulation unless otherwise stated), call duration, and call-type composition. Comparisons involving more than two stimulation conditions (e.g., baseline vs. 10, 20, 30, and 50 Hz stimulation) were analyzed using ordinary one-way ANOVA. When a significant main effect was detected, Dunnett’s multiple comparisons test was used to compare each stimulation condition to baseline while controlling the familywise error rate. In cases where the omnibus ANOVA did not reach significance, post hoc comparisons were still reported when prespecified comparisons to baseline were performed.

For developmental analyses of pup USVs across postnatal ages (P0–P21), one-way ANOVA was used to assess effects of age within genotype, with genotype comparisons performed at individual time points using unpaired two-tailed *t* tests where appropriate.

#### Call-type composition analyses

Call-type proportions were analyzed using centered log-ratio (CLR) transformation to account for the compositional nature of vocal repertoire data. For each animal, CLR differences between stimulation and baseline were computed for the first 100 calls following vocal onset. Effect sizes were summarized using paired-sample Cohen’s *d* (mean ΔCLR / SD across animals). Statistical significance across call categories was assessed using paired comparisons with false discovery rate (FDR) correction (Benjamini–Hochberg procedure). Heatmaps display mean ΔCLR values across animals. For region exploration experiments (Fig. 3A), the proportion of call types were compared to baseline and averaged across animals for each region within the 30 s stimulation window.

#### Social interaction metrics

Social interaction was quantified using SLEAP-derived pose tracking to compute Euclidean distances between animals (nose-to-nose, nose-to-tail base, and center-to-center distances, Movie S3). For optogenetic experiments, distances were averaged over defined epochs (0–30 s stimulation and 30–60 s post-stimulation). Effects of stimulation frequency were assessed using ordinary one-way ANOVA followed by Dunnett’s multiple comparisons test versus baseline. Control littermate datasets were analyzed in parallel using identical statistical procedures.

#### Tremor analysis

Tremor power spectra obtained from the accelerometer-based tremor monitor were analyzed by calculating the maximum power within the stimulation frequency band for baseline and stimulation periods. Paired two-tailed *t* tests were used to compare baseline and stimulation power for each frequency and targeting condition. For experiments involving multiple stimulation frequencies, each frequency was treated as an independent comparison relative to baseline.

#### Motor phenotype scoring

Binary presence or absence of tremor, ataxia, and dystonia was recorded by at least two expert observers blinded to stimulation condition. For dystonia severity ratings (0–5 scale), baseline and post-DBS values were compared using paired two-tailed *t* tests. Because the dystonia scale is ordinal, results are interpreted conservatively and reported alongside individual data points.

#### Real-time place preference (RTPP)

RTPP behavior was analyzed by comparing time spent in stimulated versus unstimulated zones, total distance traveled, and number of zone entries. Paired two-tailed *t* tests were used to compare light-OFF and light-ON conditions within animals. Unilateral and bilateral stimulation cohorts were pooled after confirming no qualitative differences in behavioral directionality.

#### Data presentation

Bar plots depict mean ± standard deviation unless otherwise indicated. Individual animals are represented as dots. Heatmaps represent averaged values across animals. Exact *N* values for each analysis are reported in the corresponding figure panels or legends. No data points were excluded unless predefined criteria were not met (e.g., insufficient number of USVs for call-type analyses).

**Figure S1.**
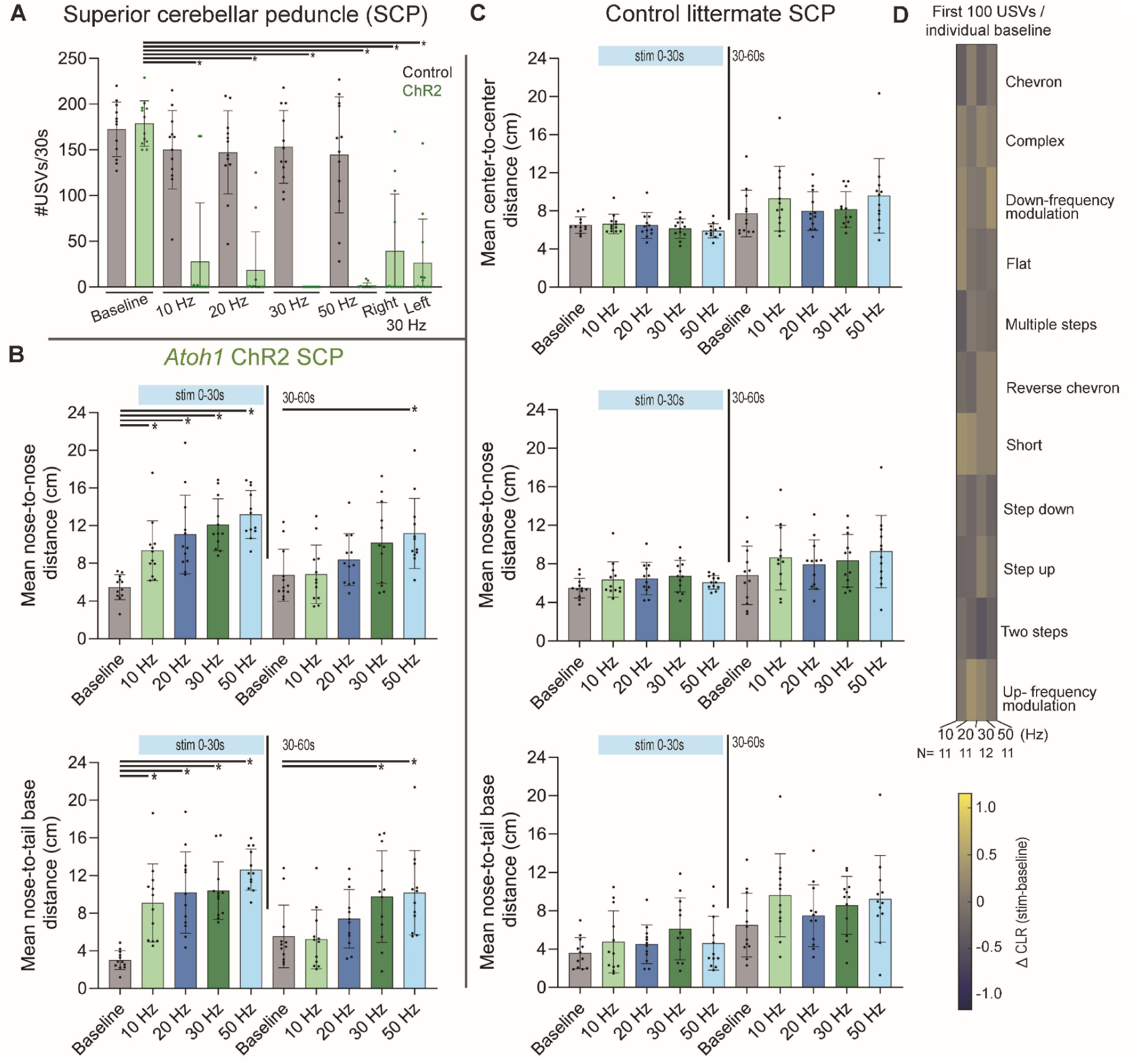
Control animals do not exhibit vocal or social suppression during superior cerebellar peduncle (SCP) stimulation. **(A)** Quantification of ultrasonic vocalizations (USVs) produced by male mice during social interaction with control littermate *Rosa26^lsl-ChR2-eYFP^* (gray) or ChR2-expressing *Atoh1^Cre^*;*Rosa26^lsl-ChR2-eYFP^* (green) animals during optogenetic stimulation of the superior cerebellar peduncle (SCP). In ChR2-expressing animals, SCP stimulation at 10–50 Hz robustly suppresses USV production relative to baseline, whereas control animals show no suppression across stimulation frequencies. One-way ANOVA revealed a significant effect of stimulation condition (*F*(6,77) = 25.64, *p* < 0.0001), followed by Dunnett’s multiple-comparisons test comparing each stimulation condition to baseline (10 Hz, 20 Hz, 30 Hz, and 50 Hz: *p* < 0.0001). Right and left unilateral 30 Hz SCP stimulation conditions are shown separately (*p* < 0.0001 for both). Error bars indicate mean ± SD; dots represent individual animals. *N* = 12 animals. **(B)** Social interaction metrics during SCP stimulation in ChR2-expressing animals. Mean nose-to-nose distance (top) and male nose-to-female tail-base distance (bottom) are shown for the first 30 s of stimulation (0–30 s; blue bar) and the subsequent 30 s interval (30–60 s). SCP stimulation significantly increases inter-animal distance in a frequency-dependent manner during both epochs, consistent with social disengagement. One-way ANOVA revealed a significant main effect of stimulation condition for nose-to-nose distance (0–30 s: *F*(4,55) = 12.57, *p* < 0.0001; 30–60 s: *F*(4,55) = 4.128, *p* = 0.0054). Dunnett’s post hoc comparisons versus baseline were 10 Hz: *p* = 0.0077; 20 Hz , 30 Hz, and 50 Hz: *p* < 0.0001. For male nose-to-female tail-base distance, one-way ANOVA revealed a significant main effect of stimulation condition (0–30 s: *F*(4,55) = 15.40, *p* < 0.0001; 30–60 s: *F*(4,55) = 4.281, *p* = 0.0044), with Dunnett’s post hoc comparisons versus baseline significant at 30 Hz (*p* = 0.0333) and 50 Hz (*p* = 0.0168). Error bars denote mean ± SD; dots represent individual animals. *N* = 12 animals. **(C)** Social interaction metrics in control littermate mice implanted above the SCP. Mean center-to-center (top), nose-to-nose (middle), and nose-to-tail-base (bottom) distances are shown for the same stimulation epochs as in (B). In contrast to ChR2-expressing animals, control mice do not exhibit significant changes in social interaction during SCP stimulation. One-way ANOVA revealed no significant effect of stimulation condition for any measure during the 0–30 s epoch (center-to-center: *F*(4,55) = 1.030, *p* = 0.403; nose-to-nose: *F*(4,55) = 1.322, *p* = 0.2732; nose-to-tail-base: *F*(4,55) = 1.388, *p* = 0.2500). Dunnett’s post hoc comparisons versus baseline were not significant. Error bars indicate mean ± SD. *N* = 12 animals. **(D)** Call-type composition during SCP stimulation in control animals. Heatmap shows the centered log-ratio (CLR) difference between stimulation and baseline for the first 100 USVs produced in each session, computed per animal and averaged across animals within each stimulation frequency (10–50 Hz). Only animals with ≥100 calls in both baseline and stimulation sessions were included (*N* = 11 per frequency). Control animals exhibit stable call-type distributions across stimulation conditions, with no systematic enrichment or depletion of specific syllable classes. Color scale denotes ΔCLR (stimulation − baseline). This contrasts with the stimulation-dependent call-type redistribution observed in ChR2-expressing animals (Fig. 1E).

**Figure S2.**
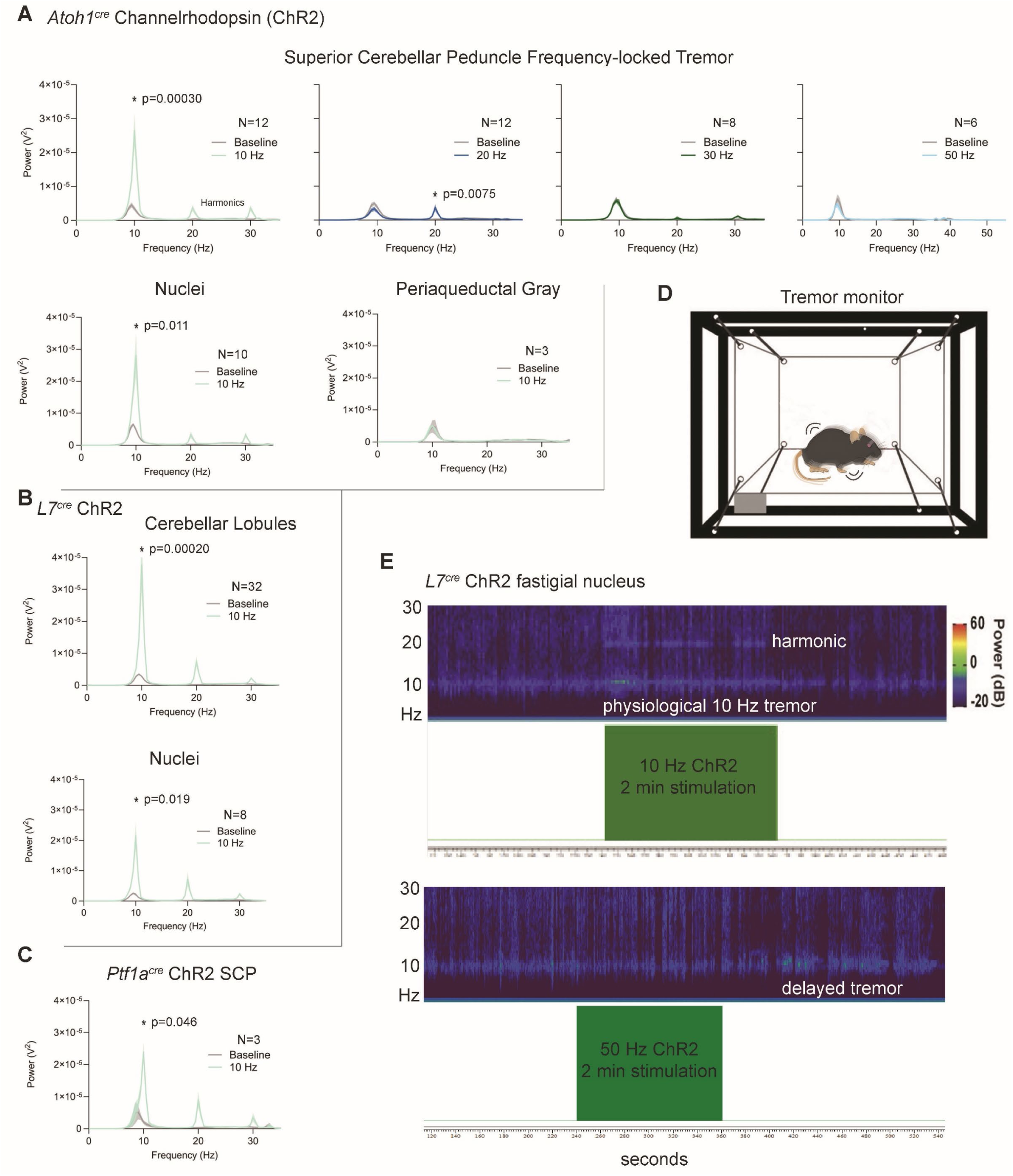
The cerebellum drives tremor across regions and cell types. (A) Optogenetic stimulation of excitatory cerebellar output neurons labeled from the *Atoh1* promoter and projecting through the superior cerebellar peduncle (SCP) generated frequency-locked tremor. Ten-hertz channelrhodopsin (ChR2) stimulation produced a prominent increase in power at 10 Hz, whereas 20 Hz stimulation increased power at 20 Hz to a lesser extent, and 30 Hz stimulation produced no significant increase at 30 Hz. Statistical significance (asterisks) was assessed by comparing the maximum power within the stimulation frequency band between baseline and stimulation using paired *t*-tests. Each stimulation was performed as an independent trial, using the 2-min pre-stimulation period as baseline. Targeting excitatory neurons within individual cerebellar nuclei (fastigial, interposed, and dentate), from which SCP projections originate, similarly increased power at 10 Hz, whereas stimulation of the same projection pathway targeting the periaqueductal gray did not elicit tremor. Tremor was identified visually in behavioral videos, and all animals within each group were visually confirmed to exhibit increased spectral power during stimulation, as illustrated in (E). (B) Ten-hertz stimulation of Purkinje cells expressing ChR2 under control of the L7 promoter increased power at 10 Hz across all cerebellar lobules examined (vermis VI, crus I, crus II, crus I/II bundle, paravermis IV/V, and vermis IV/V). A comparable increase in tremor power was observed when stimulating Purkinje cell projections to individual cerebellar nuclei. (C) Optogenetic stimulation of inhibitory cerebellar projections labeled by the *Ptf1a* promoter within the SCP also increased tremor power during 10 Hz stimulation. (D) Schematic of the tremor monitoring apparatus, consisting of a Plexiglass chamber suspended by elastic cords and coupled to an accelerometer (gray). (E) Continuous time–frequency representation (sonogram) of tremor power from a representative animal with fastigial nucleus targeting during 10 Hz and 50 Hz stimulation. Typically across trials, 10 Hz stimulation produces a 10 Hz tremor from every cerebellar region, while 50 Hz has no increase in power or visible tremor. However, when targeting the fastigial Purkinje cell projections (and interposed in other trials), a visible tremor is observed in videos. The tremor monitor showed an increase in power at 10 Hz (not 50 Hz) indicating a physiological tremor was produced, that could occur during the post-stimulation period as a “delayed tremor” (e.g. Movie S4).

**Figure S3.**
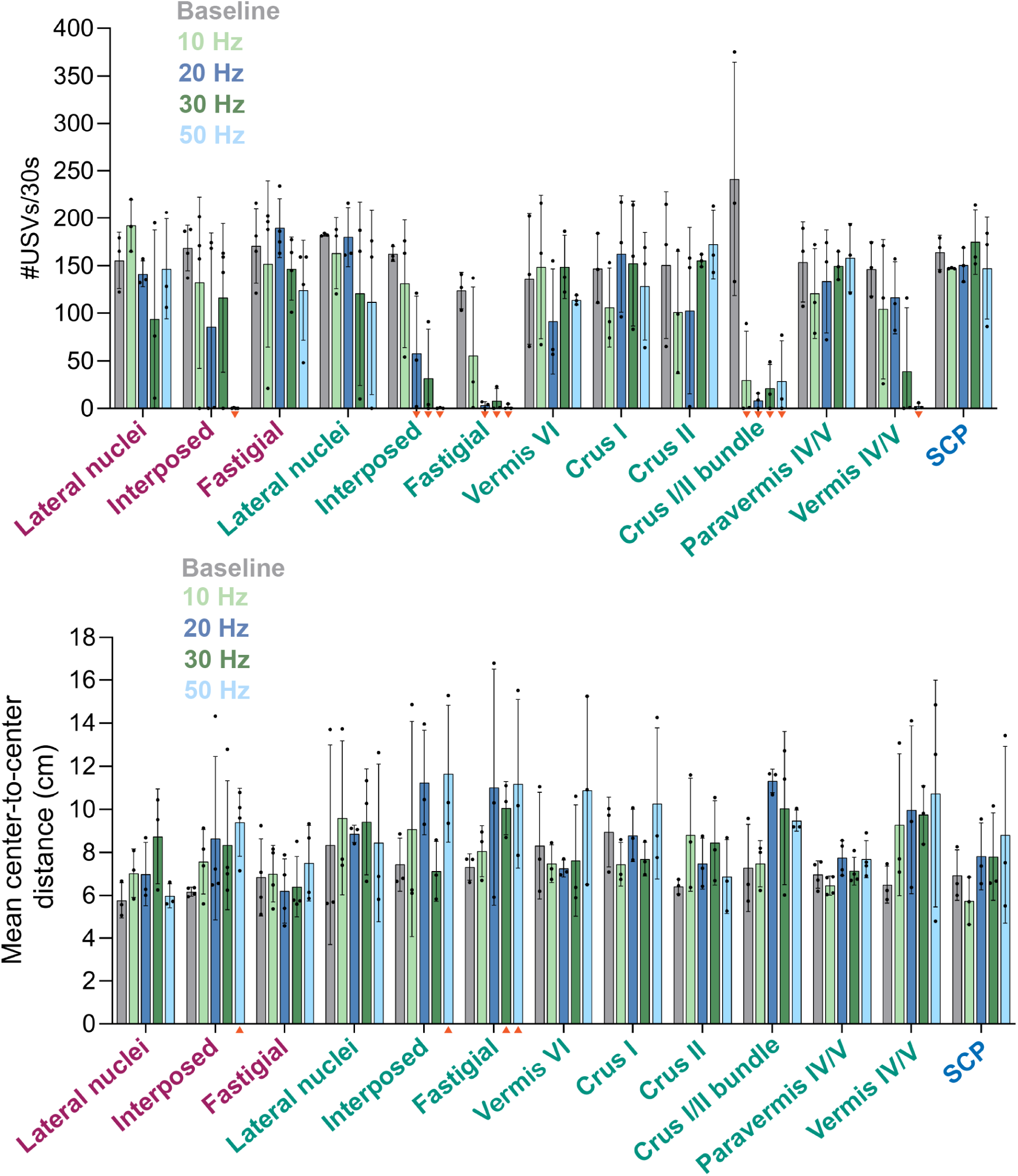
Cerebellar mapping of social communication metrics. Regions were assessed to determine which circuits and cell types alter ultrasonic vocalization (USV) production and lead to increased social distance. Due to the exploratory nature across many regions, sample sizes were *N* = 3 and we indicate trends with orange triangles shown under specific stimulation frequency. For USV number, binary phenotype was indicated as present in Fig. 2 when all three animals showed a >20% reduction in USV number. For increased inter-animal distance, a trend is indicated with an orange triangle when all animals increase distance by >12%. Excitatory SCP and PAG data are shown in Fig 1 and Fig 3, respectively.

**Figure S4.**
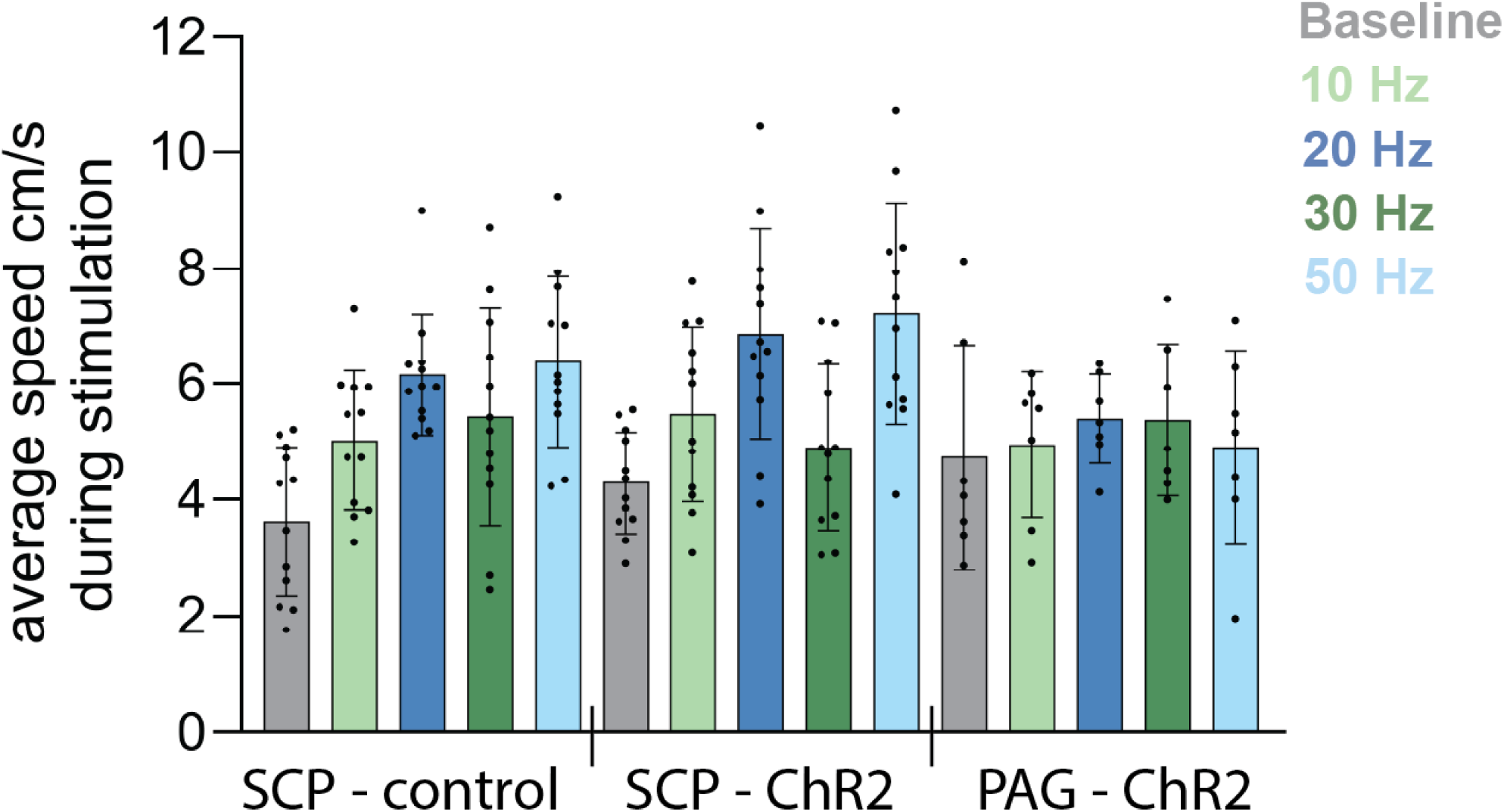
General locomotion during stimulation. Average locomotor speed measured across the 30 second stimulation period was not significantly different between control mice with bilateral SCP implants lacking channelrhodopsin expression (control: *Rosa26^lsl-ChR2-eYFP^*) and ChR2-expressing mice *Atoh1^Cre^*;*Rosa26^lsl-ChR2-eYFP^*targeting either the SCP or PAG. Thus, suppression of vocal output during stimulation was not explained by a loss of general locomotion. Movie S2 illustrates preserved locomotion despite repeated avoidance of the female, particularly during 30 Hz stimulation.

**Figure S5.**
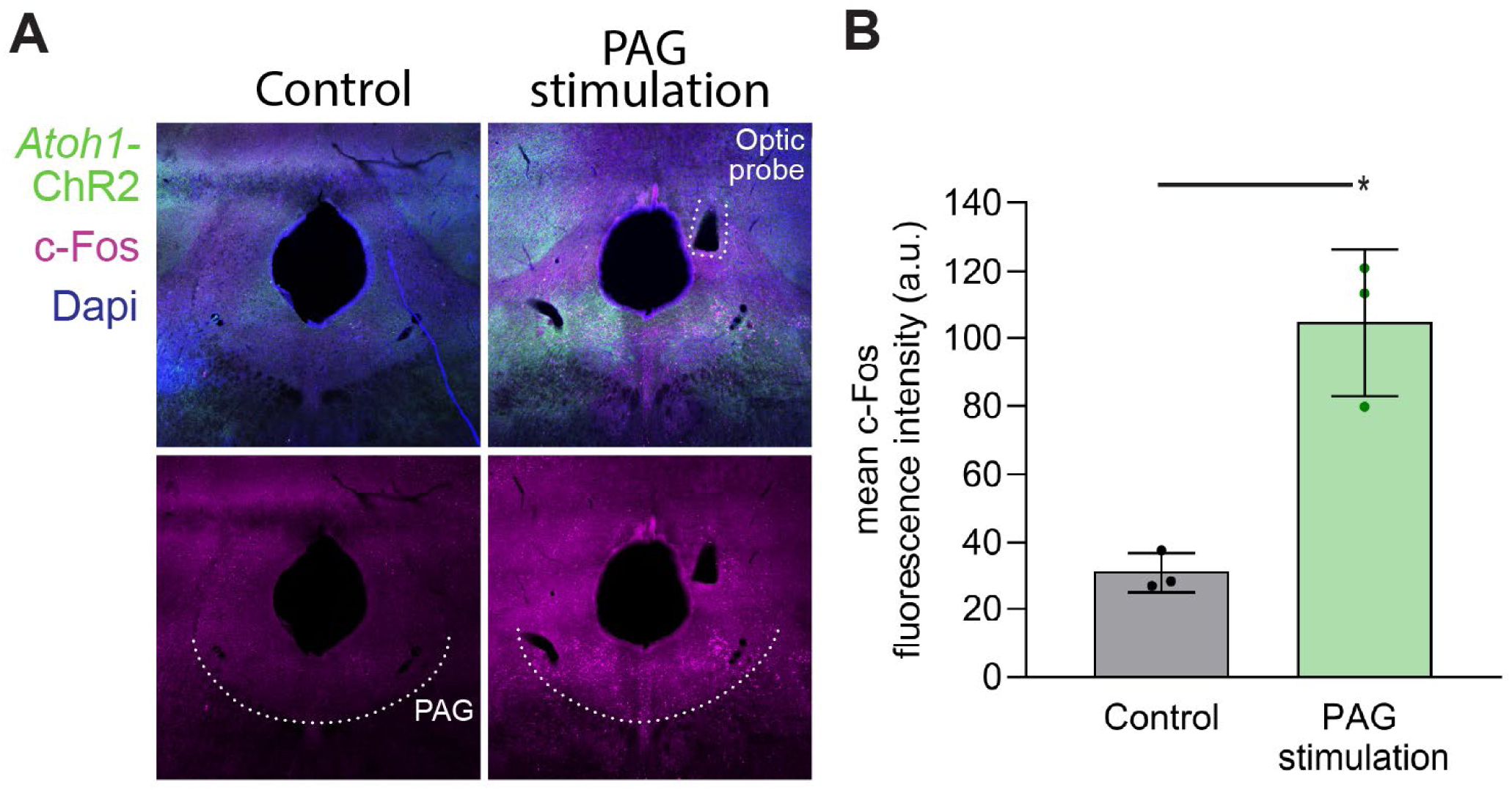
Optogenetic stimulation of cerebellar projections to the periaqueductal gray induces c-Fos expression. (A) Representative coronal sections through the midbrain showing c-Fos expression in the periaqueductal gray (PAG) of control animals (left) and following optogenetic stimulation of cerebellar excitatory projections within the PAG (right). *Atoh1*-ChR2 expression is shown in green, c-Fos immunoreactivity in magenta, and DAPI in blue. The optic probe tract is indicated in stimulated animals. Dotted lines outline the approximate boundaries of the ventral PAG. (B) Quantification of mean c-Fos fluorescence intensity within the PAG. For each animal (*n* = 3 control, *n* = 3 PAG stimulation), two anatomically matched coronal sections were analyzed, with bilateral rectangular regions of interest placed within the PAG on each section (four ROIs per animal). Mean pixel intensity values were averaged across ROIs to yield a single value per animal. PAG stimulation resulted in a significant increase in c-Fos expression compared to non-implanted controls (unpaired *t*-test, *p* = 0.0049). Bars represent mean ± SD; dots represent individual animals. (**B**) Example section of widespread activation of cells throughout the PAG. Unilateral implants showed increased c-Fos expression on both side of the PAG.

**Figure S6.**
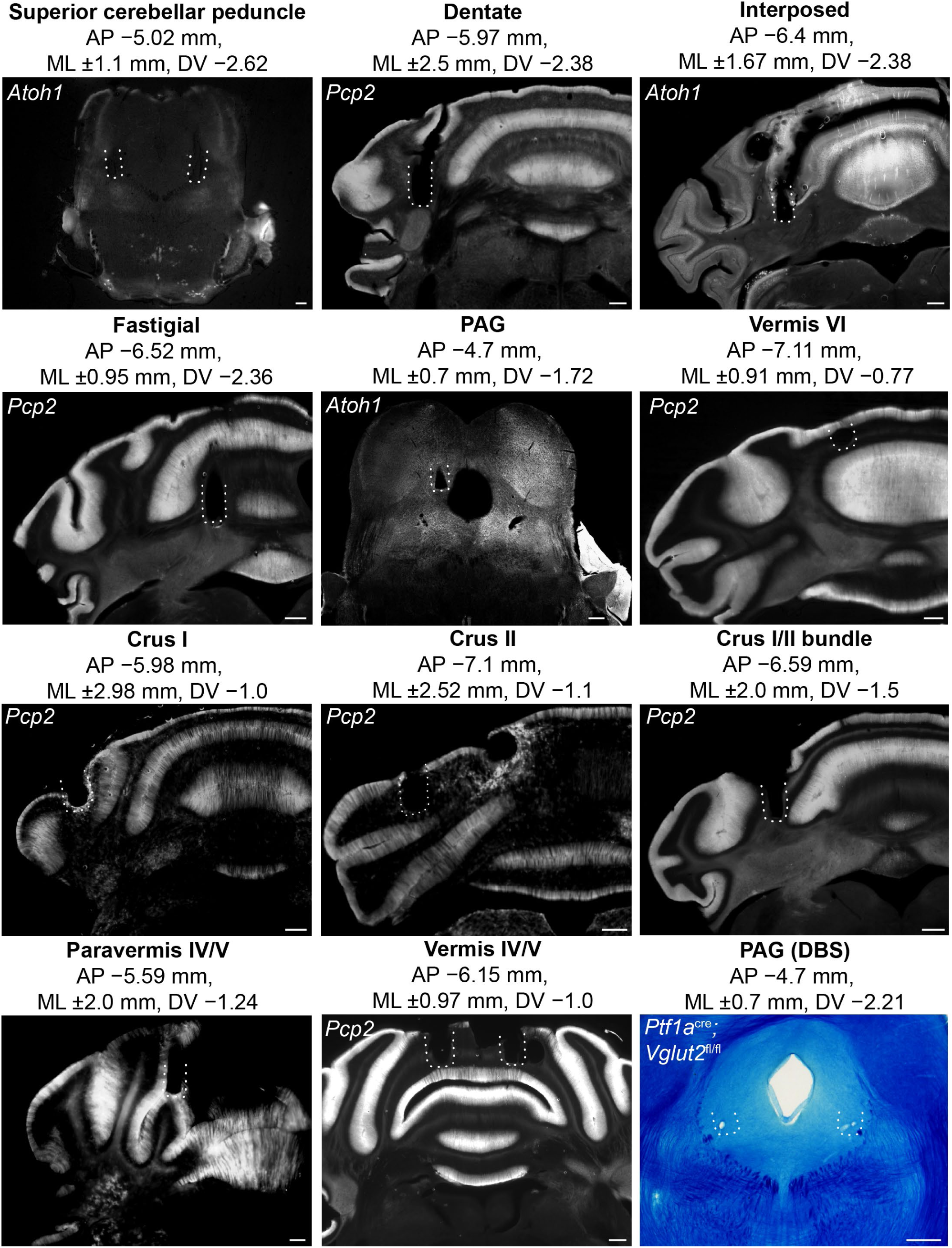
Targeted regions for optical and deep brain stimulation. Representative coronal sections from mice with surgical implants (dotted outlines) at each target location. The Cre driver line used to direct channelrhodopsin expression is indicated, along with stereotaxic coordinates relative to bregma. Fluorescence represents endogenous channelrhodopsin expression. For DBS experiments, sections were stained with Luxol Fast Blue to visualize the implant, and the electrode tip indicating the site of current delivery is shown. Scale bars, 200 µm.

**Movie S1.** 3D light sheet reconstruction showing anatomical separation of the superior cerebellar peduncle targeted for stimulation and overall Purkinje cell projection topography. Gamma was increased to display low intensity projections more prominently and the brightest green spots in the *Atoh1*-tdTomato reconstruction are not relevant, as they are brain surface artifacts (e.g. air bubbles).

**Movie S2.** Representative social interaction assays for each stimulation frequency when targeting excitatory cells within the superior cerebellar peduncle. Without stimulation (baseline), males emit vocalizations after female entry. At 10 Hz stimulation, a tremor is observed and at 20-50 Hz, increased social distance becomes visually apparent. At 50 Hz, mild motor incoordination is observed, with the male flattening low and staying still or wobbling as it crosses the chamber, sometimes extending limbs for balance and displaying brief tail oscillations.

**Movie S3.** Measuring inter-animal distance with Social LEAP Estimates Animal Poses (SLEAP) trained model. Euclidean distance between each body point displayed were measured.

**Movie S4.** Examples of behaviorally identified motor incoordination resembling tremor, ataxia and dystonia when targeting channelrhodopsin expressing Purkinje cells. Hyper-extended limbs can be seen throughout the interposed stimulation, a strong indicator of co-contraction of muscles.

**Movie S5.** Real-time place preference assay for the first 2.5 minutes in one representative animal where cerebellar excitatory projections in the periaqueductal gray were targeted with 50 Hz stimulation.

## Notes

### Competing Interest Statement

The authors have declared no competing interest.

### Summary of Updates

minor edits to figure legends minor edits to narrative phrasing

